# Atlas and developmental dynamics of mouse DNase I hypersensitive sites

**DOI:** 10.1101/2020.06.26.172718

**Authors:** Charles E. Breeze, John Lazar, Tim Mercer, Jessica Halow, Ida Washington, Kristen Lee, Sean Ibarrientos, Andres Castillo, Fidencio Neri, Eric Haugen, Eric Rynes, Alex Reynolds, Daniel Bates, Morgan Diegel, Douglas Dunn, Rajinder Kaul, Richard Sandstrom, Wouter Meuleman, M.A. Bender, Mark Groudine, John A. Stamatoyannopoulos

## Abstract

Early mammalian development is orchestrated by genome-encoded regulatory elements populated by a changing complement of regulatory factors, creating a dynamic chromatin landscape. To define the spatiotemporal organization of regulatory DNA landscapes during mouse development and maturation, we generated nucleotide-resolution DNA accessibility maps from 15 tissues sampled at 9 intervals spanning post-conception day 9.5 through early adult, and integrated these with 41 adult-stage DNase-seq profiles to create a global atlas of mouse regulatory DNA. Collectively, we delineated >1.8 million DNase I hypersensitive sites (DHSs), with the vast majority displaying temporal and tissue-selective patterning. Here we show that tissue regulatory DNA compartments show sharp embryonic-to-fetal transitions characterized by wholesale turnover of DHSs and progressive domination by a diminishing number of transcription factors. We show further that aligning mouse and human fetal development on a regulatory axis exposes disease-associated variation enriched in early intervals lacking human samples. Our results provide an expansive new resource for decoding mammalian developmental regulatory programs.

## Introduction

Mammalian *in utero* development, including organ formation and growth, is orchestrated by intricate regulatory programs that involve changes in the epigenetic landscape, the expression of transcription factors (TFs), and the functional activation of regulatory elements^1^. However, difficulties in sampling these early stages of early development have limited our understanding of these regulatory programs to date.

Early studies describing the morphological stages of mammalian development formed the foundation of developmental biology^2,3^. These foundational studies classify mouse development into pre-implantation, embryonic (days 5-11), and fetal (days 11-20) stages^4,5^. Organogenesis occurs between days 6.5 and 11 and is followed by the fetal period, which is characterized by rapid organ growth and maturation^4^. During the fetal stages, organs may also perform different roles from adult tissues. For example, the fetal liver performs hematopoietic functions^6^.

Our current understanding of the regulatory programs that govern these *in utero* stages is projected from cell line models and limited fetal samples^7,8^. While these samples suggest extensive remodeling of the regulatory landscape during *in utero* development, we still lack detailed characterization of regulatory programs for these stages *in vivo*^9^. As a result, we have an incomplete understanding of the regulatory changes, including key regulatory elements and TFs that orchestrate early mammalian development.

Understanding these early regulatory programs would not only improve our understanding of developmental biology but also provide insight into human disease etiology. A large fraction of human adult disease burden is thought to originate during early development^10,11^. However, these early developmental stages cannot be ethically sampled in humans, and are largely inaccessible for study. This challenge can be addressed, in part, by a comparative analysis of early development in the mouse, which serves as a primary experimental model for human biology^12,13^.

Here, we have used DNase-seq to profile the regulatory landscape across the late embryonic and fetal stages of mouse development. This approach maps the accessible chromatin landscape and identifies regulatory elements across different organs and developmental stages. Analysis of these datasets allows us to identify the action of key transcription factors and define the regulatory programs that govern the establishment and growth of major organs. A comparative study of these maps also reveals common principles of developmental regulation and provides insights into the origin of human disease in early development. Together, these detailed regulatory maps constitute a valuable addition to our understanding of mammalian development^2,3^.

## Results

### Regulatory DNA landscapes of mouse development

To map regulatory DNA landscapes spanning mouse development, we generated DNase-seq profiles for a wide range of samples spanning embryonic, fetal, and adult mouse stages and all major organs and tissues (**Figure 1A**)^14^. We extensively sampled tissues within different embryonic and fetal time-points (**Supplementary Table 1**), vastly extending the previous scope of accessible DNA samples for these stages. We applied strict quality control, including filtering for signal-to-noise ratios^15^, DNA fragment size distribution, and library complexity metrics, resulting in reference-grade datasets that were aggregated with other datasets from the ENCODE project^16^, yielding a total of 197 mouse samples. Samples spanned 15 developing and neonatal mouse tissues, including forebrain (7), midbrain (6) and hindbrain (8), lung (6), heart (6), thymus (4), kidney (4), limb (6), stomach (2), craniofacial tissue (5), retina (5), Müller retinal glial cells (3), and neural tube (3), among others **(Figure 1A, Methods)**. We generated tissue DNase-seq profiles within the scope of 9 time-points (post-conception days 9.5, 10, 10.5, 11, 11.5, 14.5, natal day P0, postnatal day 7, and adult stages).

**Figure 1:**
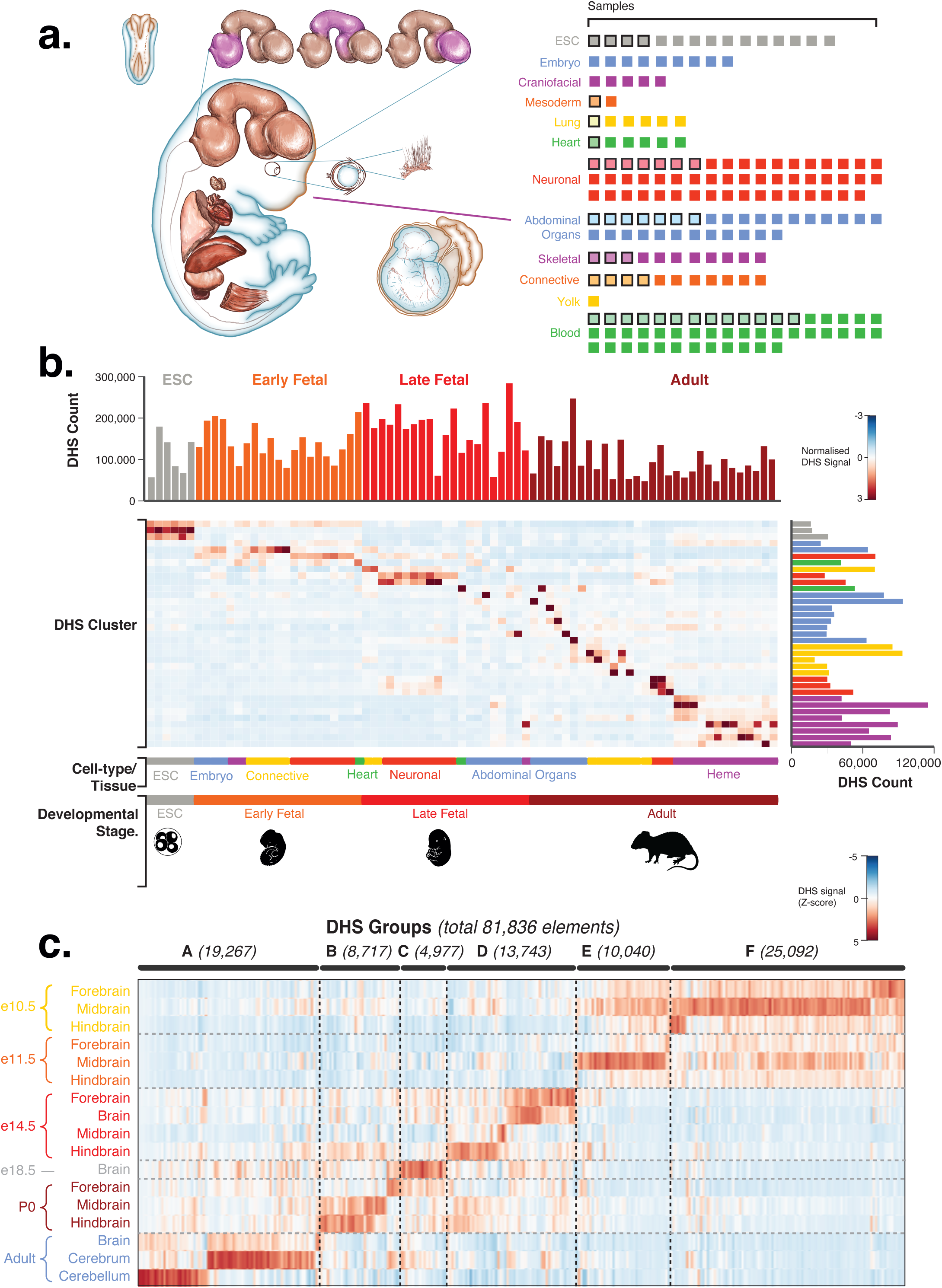
An atlas of mouse development across tissues and time-points. **[A]** List of fetal tissues analyzed with DNase-seq. Fetal time-points include days 9.5, 10, 10.5, 11, 11.5 and 14.5. Each box in the diagram on the right corresponds to a single analyzed sample. Boxes in bold correspond to samples from Vierstra et al., 2014. All other boxes correspond to new samples included in this study **[B]** K-means clustering of mouse DNase-seq data showing mouse accessible chromatin variation across time-points and tissues for 1.8 million DHSs. Primary column ordering is by stage. Secondary ordering is by tissue. Horizontal and vertical bar plots indicate the number of DHSs for each cluster and for each consolidated sample, respectively. **[C]** Heatmap of DNaseI accessibility for > 80,000 DHSs identified as differential across mouse brain development (F-test, Benjamini Hochberg FDR < 0.05). DHSs are ordered and grouped by the time-point of their maximal activity.

We performed DNase-seq at an average depth of 160.87 million uniquely mapped reads per sample. We used the hotspot2 algorithm to compute DNase I hypersensitive sites from these data (DHSs, **Methods**), identifying an average of 124,487 DHSs per sample. Among primary samples, we observed the highest number of DHSs in samples of early fetal brain tissue (321,338), and the lowest (34,693) in CD4+ T cells, a highly specialized cell type. From a cumulative total of 24,524,056 DHSs in individual samples, we merged all tissues and developmental stage/time samples to create a comprehensive atlas comprising 1,802,603 distinct DHSs, each of which was identified independently in one or more samples using the Index pipeline (**Methods, Supplementary File 1**)^17^. Within the atlas, over 467,900 elements (26%) were unique for developing or neonatal samples and were not found in DHSs from 28 adult tissues^1,16^, revealing a large untapped regulatory compartment within developmental stages.

In agreement with previous findings, the global regulatory DNA landscape of mouse development defined by DHSs was highly dynamic^9^. Only 2% of DHSs were persistently active (present in >75% of samples) across tissues and time-points, of which 46% were promoters of protein-coding genes, consistent with prior observations concerning the tissue promiscuity of promoters^18^. The vast majority of DHSs were selective for an individual tissue (32%), a particular developmental stage (20%), or an organ system of functionally related samples (46%, **Supplementary Figure 1B**); >86% were located in intronic and intergenic compartments >2.5kb from the transcriptional start sites of annotated protein-coding genes.

To gain a global view of variation in regulatory DNA profiles across all samples, we performed k-means clustering of DHSs by signal intensity (**Methods**). This analysis enabled classification of tissue-selective DHSs as a function of developmental stage (**Figure 1B,** consolidated samples listed in **Supplementary Table 2**), indicating specialized roles of the constituent elements in governing specific developmental programs as part of consistent stage and tissue-specific clusters. K-means clustering revealed extensive organ- and time-point-specificity for mapped DHSs, with the largest clusters found for adult immune cell samples, adult connective tissue, and late fetal abdominal organs. Interestingly, a large proportion of clusters (10/35, 28.57%) were localized to specific tissues and organs for the early and late fetal period, which is indicative of a large proportion of previously unmapped genomic regulation restricted to these particular stages. Importantly, this temporal patterning indicates mapping of specific regulatory programs that regulate the scheduled transition from early development and growth to later functional stages.

To further understand the widespread variation in regulatory DNA profiles for a particular organ, we sought to characterize the dynamics of temporal DHS variation across one of our most extensively assayed tissues. Specifically, we focused on the developing brain, which was the most comprehensively sampled organ. Between post-conception day 10.5 (e10.5) and day of birth (postnatal day 0 or P0), we identified 81,836 DHSs from forebrain, midbrain, and hindbrain that either (i) arose *de novo*, or (ii) were present at an earlier time-point but were then completely extinguished, or (iii) underwent a significant amplification or diminution of DNase-signal (**Figure 1C, Supplementary Figure 4B, Methods**). These findings further confirm the presence of extensive time-point- and regional-specific variation within a particular organ, highlighting the mapping of regulatory DNA elements underlying programs governing brain regional variation.

### Measuring transcription factor activity during fetal development

To understand the regulatory programs underlying DHS profiles, we incorporated TF motif and promoter signal data to measure activity for individual TF genes (**Methods**). The proportion of DHSs overlapping known TF motifs, a metric termed TF motif accessibility, is known to provide a robust measure of transcription factor activity^16,18^. We calculated TF motif accessibility for each sample, and we then Z-score transformed this metric to enable accurate comparisons between samples and TF motifs (see **Methods**). Our analysis successfully highlights the activity of motifs belonging to many known lineage-specifying TFs in different tissues, including *Tbx20* in heart, *Otx2* in the retina, and *Neurod2* in brain development (**Supplementary Figure 2A**). To this metric, we then incorporated differences in TF promoter accessibility across samples to distinguish activity for individual TFs, as many TF motifs are non-specific (**Methods**).

Promoter accessibility is known to correlate highly with gene expression values^19,20^. We confirmed this in our data by comparing DHS intensity at promoters (TSS +/− 5kb) with RNA-seq expression values for all protein-coding genes across our samples, returning a mean correlation of r=0.69 (**Supplementary Figure 2B, C**). We used this approach to identify key lineage-specifying regulators, such as *Mef2c* during heart development or *Nkx2-1* in lung development^21,22^ (**Supplementary Figure 2D, E**). Our analysis highlighted the importance of considering both TF motif and promoter information. For example, while DHSs harboring the family of *Nkx* motifs are highly accessible in fetal lung samples, by examining promoter strength we can specifically identify *Nkx2-1* as active in fetal lung. Considering both TF motif accessibility and promoter accessibility also anticipates novel roles for TFs in mouse development. For example, we found that *Bhlhe23*, which remains to be characterized in brain formation, was specifically active during midbrain development (supported by a Z-score > 1 and 2.80-fold increase in motif accessibility, and a 1.63-fold increase promoter accessibility between e10.5 and e14.5; **Supplementary Figure 4A**).

In addition to distinguishing between TF family members, a comparison of motif accessibility and promoter accessibility can also indicate the mechanism of TF action. For most TFs (79.3%), promoter accessibility and motif accessibility are positively correlated, suggesting increased activity of the TF increases chromatin accessibility. However, for a minority of TFs that appear to condense chromatin, promoter accessibility and motif accessibility are negatively correlated, including cases of known repressive TFs such as *Gfi1b*^23^, *Snai2*^24^, and *Zbtb16*^25^ (**Supplementary Figure 3A-D**).

### Sharp turnover of regulatory programs partitions in utero development

The extensive shift in DHS numbers we observed across all samples in the k-means analysis and the fetal brain DHS analysis (**Figure 1B, C**) was accompanied by a dramatic turnover in TF motif accessibility (**Figure 2A**), TF motif accessibility correlation (**Figure 2B**), and the daily rate of change in the TF landscape (**Figure 2C, D**) occurring between e11.5 and later time-points.

**Figure 2:**
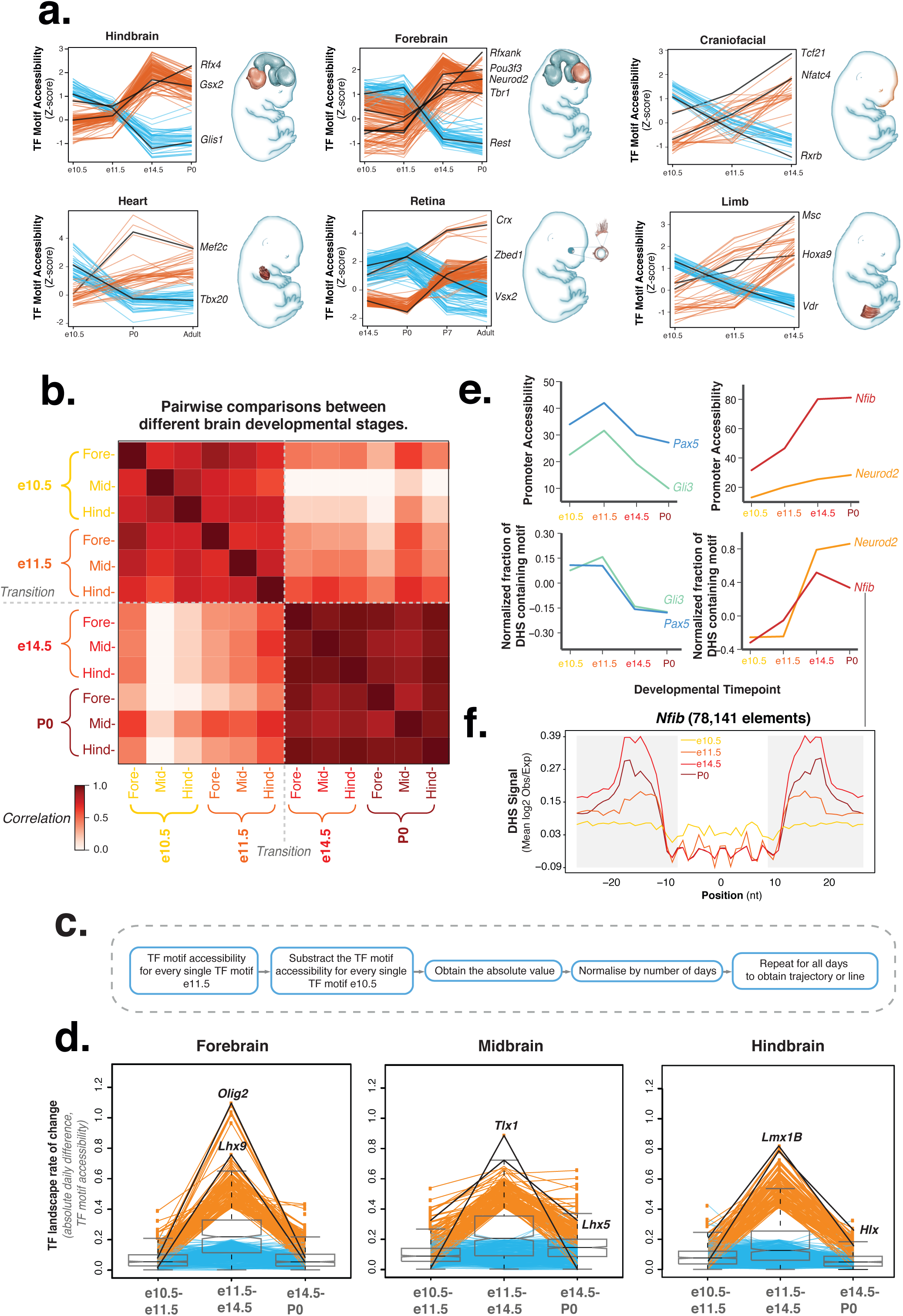
Early and late mouse regulatory programs. **[A]** Linegraphs of TF motif density with the highest variability during development for a range of mouse tissues. TFs are colored based on whether they show increasing or decreasing motif density. **[B]** Correlation in motif density in DHSs across developing brain samples. Notable is the sharp split between early and late fetal samples. **[C]** Schematic of calculation of the TF landscape rate of change or absolute daily difference in TF motif accessibility. **[D]** Daily rate of change in the TF landscape, computed as the average of the absolute value of the difference in motif density between every motif, slows down by e14.5, identifying little change between e14.5 and P0 compared to e11.5-e14.5. Line plots depict all TFs with an e11.5-e14.5 rate of change of over 0.4 (orange) and below 0.2 (blue). **[E]** Expression and DHS motif density for TFs identified as variable across developmental time, including early regulators (*Pax5*, *Gli3*), which present reduced activity over time, and neural-specifying TFs (*Nfib*, *Neurod2*), which present increased activity over time. **[F]** Aggregate DNaseI cleavage signal at the motifs of *Nfib*, a TF identified as variable across developmental time.

To evaluate the shift in the activity of individual TFs at this transition event, we computed TF motifs with significantly different Z-scores between sampled stages (**Methods**). This analysis identified 343 (54% of total) TF motifs that show differential accessibility across brain development (**Supplementary Figure 5**). For example, motifs for embryonic TFs, such as *Pax* family members and homeobox TFs, undergo condensation and repression while neural-specifying TF motifs, such as *Nfib* and *Neurod2*, become activated between e11.5 and e14.5 (**Figure 2E, F**). In the case of *Nfib,* promoter activity increases 1.7-fold with a concomitant 1.76-fold increase in its genome-wide binding levels between days e11.5 and e14.5 (**Figure 2E, F**).

We next analyzed DNase-seq signal across other developing mouse tissues for evidence of a similar regulatory transition between embryonic and fetal stages of development. For these tissues, we identified the largest turnover of regulatory elements and TF motif accessibility to occur between e11.5 and e14.5 (**Figure 2A** –craniofacial and limb panels–**, Supplementary Figure 4A, B**), in agreement with our findings in brain tissue. We found an average correlation of DNase-seq data between e11.5 and e14.5 samples of 0.59 (compared to an average correlation of 0.70 between early developmental stages and 0.79 between late fetal stages). In addition, simultaneous changes in TF gene promoter accessibility also support this transition (average correlation of 0.90 and 0.92 between e10.5/e11.5 and e14.5/P0 compared to an average correlation of 0.71 between e11.5 and e14.5).

This major regulatory transition coincides with the switch from the initial patterning and formation of organs during embryogenesis to their subsequent growth and maturation during fetal stages of development. We detected changes in regulatory programs across tissues reflecting this transition. For example, we observe the replacement of TFs that regulate heart tube formation and folding, such as *Pitx2*^19^, *Eomes*^20^, and *Six1*^21^, by TFs that specify cardiomyocyte identity such as *Mef2c*^11^, *Tbx20*^15^, and *Smad6*^22^ (**Supplementary Figure 6A, B**). Therefore, associated regulatory patterns suggest that this major developmental transition involves the clearance of early regulatory programs that govern the establishment of the body plan, which are then replaced by regulatory programs that govern the growth and maturation of individual organs.

In addition to detecting a strong developmental shift between embryonic and fetal programs, we also observe that global TF activity patterns become increasingly dominated by a small number of TFs in late developmental stages (centralized domination, **Figure 3A**). To quantify how the regulatory activity in tissues becomes increasingly dominated by a small number of cell-specific TFs, we applied the Gini index – a measure of economic inequality – to the accessibility around all TFs to measure inequality in TF activity for each developmental stage. We found that the Gini coefficient increased with development, highlighting an increasingly unequal distribution of TF activity (*P*=5.11e-08, **Figure 3B, C**). This phenomenon is conserved across tissues, with notable examples in heart, midbrain, and craniofacial tissue development (**Figures S6C-H**). These findings suggest a regulatory principle shared across tissues whereby the potential for TF activation becomes increasingly restricted to a few key TFs as development advances and tissue identity becomes entrenched.

**Figure 3:**
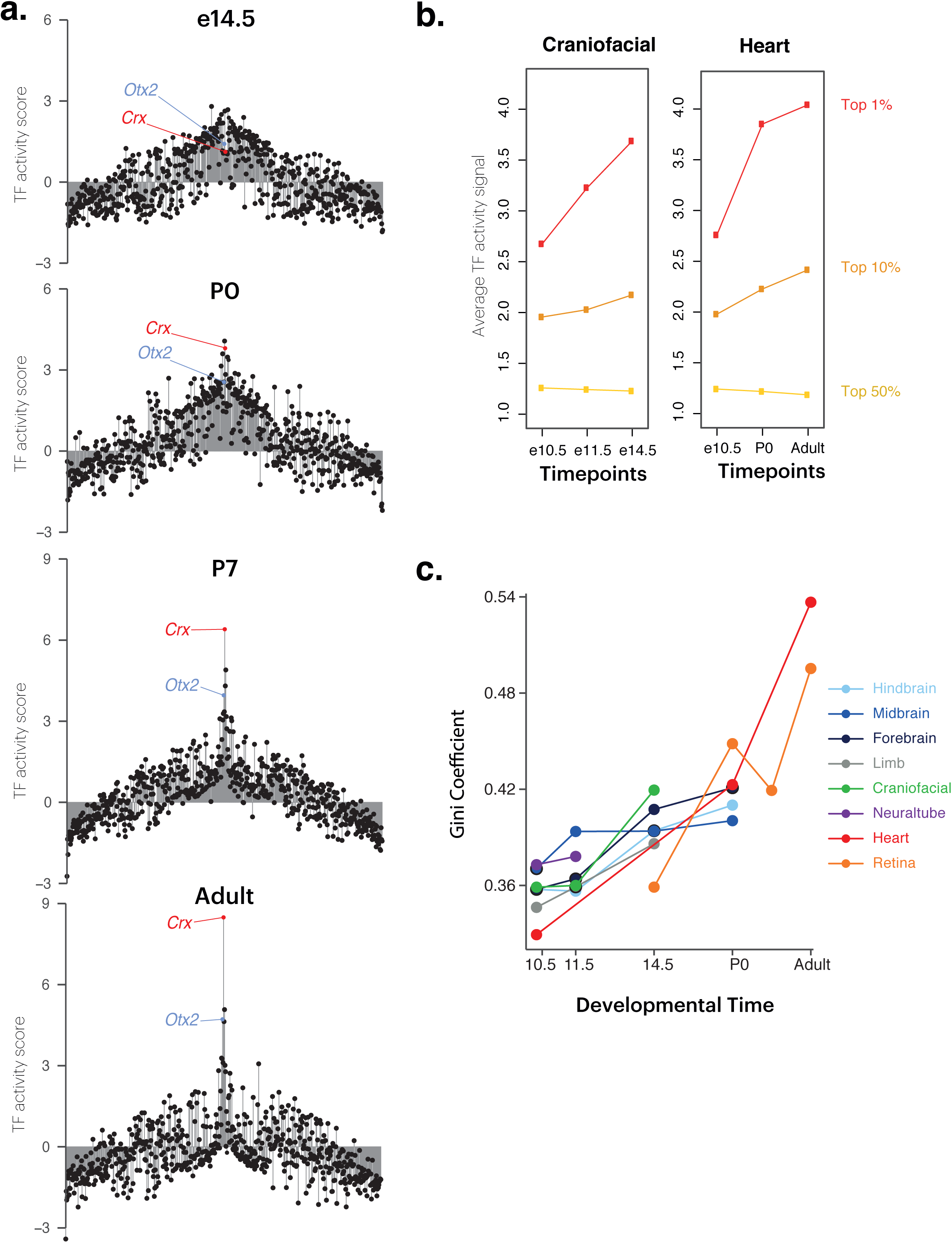
Dynamics of TF inequality across developmental stages. **[A]** Key TFs dominate retina development (centralized domination). Shown are dynamics of TF activity (combined promoter/motif score) across retina development. The height of the bar displays the TF’s activity (combined score) in each sample. The 638 TFs are arranged on the x-axis in the same manner in each Figure with the center-out ordering of the TFs determined by the average combined score across all retina samples. *Crx* and *Otx2* are known regulators of photoreceptor cells. **[B]** Average TF promoter DNase signal (normalized, +/− 5kb from TSS) for the top 1%, top 10% and top 50% of TFs at each time-point for craniofacial tissue and heart. **[C]** Gini coefficient of DNaseI cleavages in DHSs around the TSSs (+/− 5kb) of TFs across different tissues in mouse development.

### Aligning mouse and human development on a regulatory axis

We next focused on the relationship between mouse and human regulatory programs. The cis-regulatory landscape is known to present large differences at a local level between mouse and human, with an evolutionary turnover of over two thirds of the cis-regulatory landscape (**Supplementary Figure 7**)^16^. However, despite this fact, higher-level features of transcription factor-driven regulatory programs present stable conservation between human and mouse cognate organs and tissues, including global normalized patterns of TF recognition site accessibility^16^ and TF regulatory networks^26^. For virtually every human and mouse TF, the global normalized density of recognition sequences within the accessible DNA of a particular tissue (e.g., immune cells, brain, etc.) is strikingly conserved between human and mouse (**Fig. 4A**). Given this high-level conservation, we sought to verify if, given comparable data, it would be possible to create an empirical ‘alignment’ of human and mouse development based on global regulatory features. Such an alignment would be a useful and orthogonal complement to classical models such as Theiler stages in mouse^2^ and Carnegie stages in human^3^, which can be broadly aligned based on morphological criteria yet progressively diverge as development progresses.

**Figure 4:**
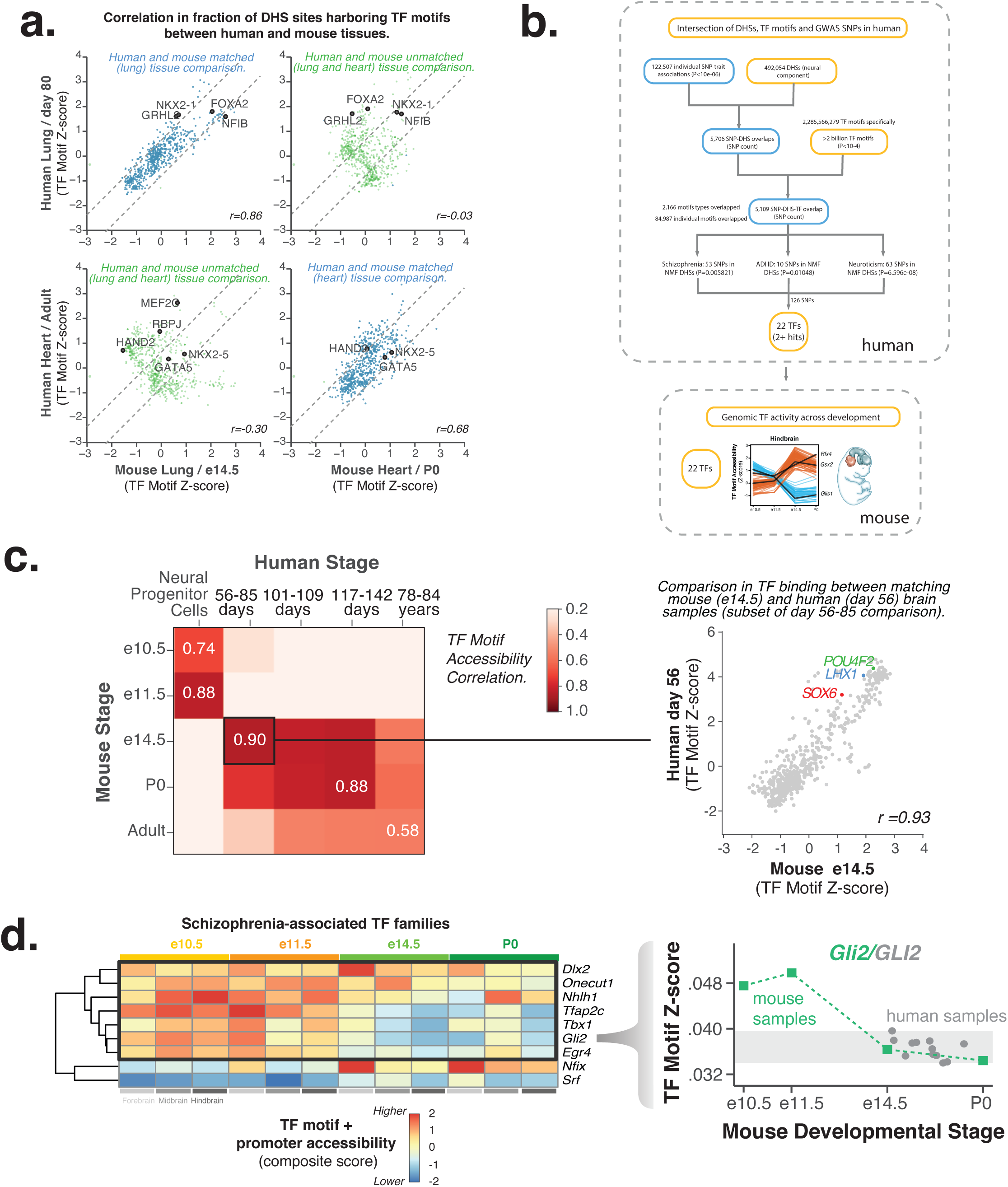
TF motif accessibility aligns human and mouse development. **[A]** Correlation of TF activity (z-score transformed motif density in DHSs) between human and mouse heart and lung samples shows the highest value between the same tissues for the two species (also shown in Vierstra et al., 2014). Given this high correlation between human and mouse genome-wide TF activity at similar stages and tissues, mouse DNase-seq landscapes can be used as a tool to study TF activity levels in early human development. **[B]** Flowchart of the analysis procedure to obtain GWAS-associated TF families. Starting with all unique SNP-trait associations present in the NHGRI-EBI GWAS catalog, we filtered for overlap with neural component DHSs (obtained as described in Meuleman et al., 2019) and TF motifs. This analysis highlighted several neural traits and TF motif families. We then evaluated the activity of TFs within these families across mouse development. **[C]** Correlation of TF activity (z-score transformed motif density in DHSs) between human brain and mouse forebrain development. To the right is an example correlation chart between one mouse and one human sample. **[D]** Early GWAS signal observed for TFs belonging to schizophrenia-associated TF families. Time-points sampled in mouse development shown above. Several TFs with motifs overlapping schizophrenia SNPs (in human) present a high brain activity signal within early mouse samples (shown are z-scores of TF genome-wide binding as heatmaps). To the right is a line plot showing *Gli2* TF motif Z-score in mouse (colored) and for *GLI2* in later human samples (grey). TF motif Z-score values for early mouse point to a higher activity for this GWAS-associated TF family in early human development.

We thus computed the normalized global density of recognition sites for each TF within DHSs in 12 human brain samples spanning the post-conception days 56 through 150 and then correlated these values with the same metric derived from DHS profiles of staged mouse forebrain samples. Using this approach, we found that day 56 human brain samples align markedly better with mouse e14.5 forebrain (Pearson’s r = 0.88) compared with other forebrain time-points (Pearson’s r = 0.43, **Figure 4B**). This temporal alignment is in precise agreement with that derived from detailed comparative morphological analyses^25^. Extending this approach to align later mouse forebrain developmental stages with human developmental samples revealed that human brain samples from days 117-142 correspond most closely with P0 mouse forebrain (Pearson’s r = 0.83-0.92). These results provide an alignment that cannot be achieved by classical models due to divergent morphology (**Figure 4B, Supplementary Figure 8A**).

We next explored the degree to which mouse development could act as a surrogate for early human developmental stages that cannot otherwise be accessed, by extending our approach to the full range of available mouse developmental and human prenatal samples. For very early time-points in brain, by morphology alone, days e10.5-e11.5 of mouse brain development are predicted to correspond with human post-conception days 20-36. As human primary samples are lacking from this period, no comparable human brain sample can be found for comparison with the early mouse samples. Instead, human neural progenitor cells (NPCs) provided the highest correlation for any human samples with the pre-e14.5 mouse brain samples, showing that early mouse samples correspond to an early unmapped human neural stage (R = 0.74, **Figure 4B, Supplementary Figure 8A**). Importantly, for human NPCs, the magnitude of the correlation is reduced compared to the temporally matched primary samples. Since the early mouse samples represent time-points unexamined in human primary datasets, we can use these to model TF activity in early human development. We thus used a linear mixed model (**Methods**) to estimate, for each TF, the density of its recognition sites in accessible DNA at human days that are otherwise unavailable to experimental interrogation (**Supplementary Figure 8B, C**).

### Using mouse development to illuminate human disease-associated variation

Genetic susceptibility to many adult-onset human diseases is thought to be linked to early developmental events^10,11^. In addition, common variants associated with human neuropsychiatric disorders are enriched in regulatory DNA active within human brain-derived samples^7,27,28,29^. However, whether these variants impact early regulatory programs cannot be determined systematically using early human samples (prior to post-conception day 56) due to a combination of ethical considerations, impracticality, and societal norms.

Our results from the mouse-human temporal alignment analysis (**Figure 4B**) indicate that samples from mouse embryonic day 11.5 and earlier correspond to samples prior to post-conception day 56 of human development. This finding, taken together with the conservation of global regulatory programs between human and mouse^16^, indicates that early mouse regulatory programs constitute a robust model for early human regulatory patterns that cannot be mapped experimentally. We thus sought to verify whether regulatory programs targeted by variants associated with human neuropsychiatric diseases were active in day 11.5 and earlier mouse samples.

To address this, we first identified all human variants for each phenotype within the NHGRI GWAS catalog^30^ and intersected these with human neural component DHSs from the ENCODE project^17^. As anticipated and previously described^7,27,28,29^, significantly enriched phenotypes included neurological traits and neuropsychiatric disorders comprising, among others, schizophrenia, neuroticism, and attention deficit hyperactivity disorder (**Figure 4C**). For these 3 disorders, we then identified the subset of the 126 unique DHS-localizing variants that could be resolved to well-annotated transcription factor recognition sites within these DHSs. We then catalogued the impacted TF motifs (**Supplementary Table 3**). Many of these TF motifs belong to families in which members have either differing recognition site preferences or, more commonly, have similar recognition sites yet present discordant tissue or temporal expression patterns. To employ a more specific metric of TF activity in mouse brain samples at a given time-point, we applied the aforementioned combined score of normalized DNase I accessibility at the promoter and motifs of each TF (**Methods**).

This analysis revealed that that TF-driven regulatory programs associated with variants linked to human neuropsychiatric disorders include TF families that show early developmental patterns of activity in mouse samples (**Figure 4C**). For example, the majority of TF families (77.7%) with recognition sites impacted by schizophrenia-associated variation are more active in early vs. late mouse brain development (**Figure 4C, Supplementary Figure 9A**). Additional early TF families were observed for other neuropsychiatric disorders **(Figure 4C, Supplementary Figure 9B-F)**.

Taken together, these results indicate that human and mouse developmental timing can be grossly aligned on a regulatory axis. This alignment can then be applied to analyze key features of disease-associated regulatory programs that are not accessible experimentally in early human samples^16^.

## Discussion

This study presents the most detailed and comprehensive survey of genome regulation during *in utero* mouse development to date. Regulation during development is both dynamic and specific, as evidenced by the large fraction of elements that are uniquely identified within this study and not described elsewhere.

Detailed analysis of these DNase-seq maps allows us to trace the regulatory programs that define each organ and tissue throughout mouse development. Further analysis enables the identification of the key transcription factors and regulatory elements that drive these developmental programs. Within this study, we have highlighted a few notable examples. However, we provide the DNase-seq maps as a detailed resource through which biologists can further analyze the developmental regulation of individual tissues and organs.

The breadth of profiled organs, tissues, and stages also enables global events and principles to be identified. Most notably, our study showed that the regulatory landscape undergoes a major transition between embryonic and fetal stages of development. This transition, which complements observations from anatomical studies, involves a general turnover of regulatory programs, with the clearance of earlier embryonic programs of tissue patterning and formation, which are replaced by programs that govern organ growth and function. Following this transition, the regulatory landscape becomes increasingly unequal as it is increasingly dominated by a few high-activity lineage-specifying transcription factors that enable tissue identity and growth to become entrenched within ongoing development. As a result of both the transition and the rising inequality in TF activity, we found the regulatory landscape of late developmental stages to be markedly different from that of early stages.

Our profile of mouse developmental regulation can also supplement gaps in our understanding of human development. Due to ethical considerations, many analogous stages in human development cannot be sampled, and we rely on cell line studies that poorly recapitulate the complex morphology, context, and environment under which development occurs. Accordingly, the profile of mouse developmental regulation can complement anatomical studies and supplement our incomplete understanding of early human development. This understanding is becoming increasingly important, following a growing appreciation of the role of early development in the origin of specific complex diseases, such as schizophrenia.

Early studies describing morphological stages during ontogeny established the field of developmental biology^3^. This canon of studies produced a framework for understanding the procession of stages by which an organism develops. Here, we complement those early morphological studies with detailed and comprehensive regulatory maps for these same developmental stages. We anticipate that, like those early studies, this resource will provide a comprehensive framework in which to analyze and understand the intricate regulatory programs that govern mammalian development.

## Supporting information

File S1: Mouse Atlas BED file (197 samples, mm10)

File S2: ENCODE DCC library IDs

## Author contributions

JAS conceived the project. CEB, JL, TM, EH, ER, AR, RS, WM and JAS performed or supported data analysis. CEB, JH, IW, KL, SI, AC, FN, DB, MD, DD, RK, MAB, MG and JAS performed or supervised data collection and DNase-seq experiments. JAS supervised the project. CEB and JAS wrote the manuscript with contributions from the other authors.

## ACKNOWLEDGEMENTS

This work was supported by NIH grant 3U54HG007010 to J.A.S., and by a charitable contribution from GlaxoSmithKline. We acknowledge the ENCODE DCC and the Wold lab for providing RNA-seq data used for analyses. This work is part of the mouse ENCODE project (https://www.encodeproject.org/).

