## Supplemental Figures for "Atlas and developmental dynamics of mouse DNase I hypersensitive sites"

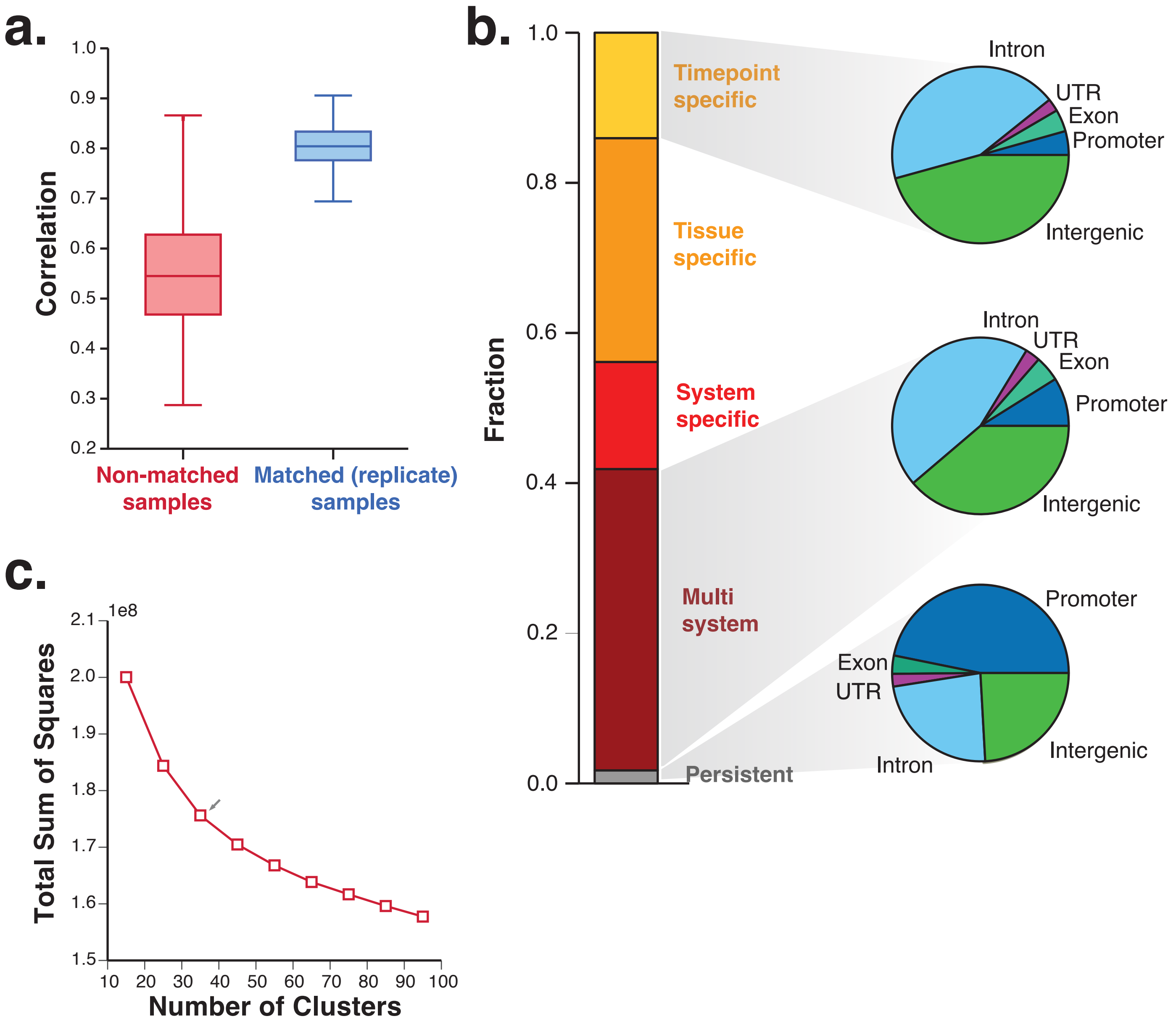


**Figure S1. Correlation and specificity between DHS libraries. (A)** Box (interquartile range, IQR) and whisker (1.5*IQR) plot of replicate concordance (Pearson correlation between replicates of log DNaseI cleavage density + 1) with unmatched samples included for comparison. **(B)** Stacked barplot showing the fraction of DHSs that are persistent (>75% of samples), tissue-specific, timepoint-specific, system-specific and multi-system. Samples were divided into Brain, Carcinoma, Connective, Cardiovascular, Gastrointestinal, Hematological, Kidney, Lung, Muscular, Stem cell, and Yolk systems. For each category of DHSs, the overlap with gene annotations is also shown (indicated with pie charts, including intron, UTR, exon, promoter and intergenic categories). **(C)** Elbow test indicating optimal number of clusters for k-means clustering (k=35). Y axis indicates the total sum of squares for each clustering model (values are shown on 1e8 scale). X axis indicates the number of clusters (or k) for each k-means clustering model.

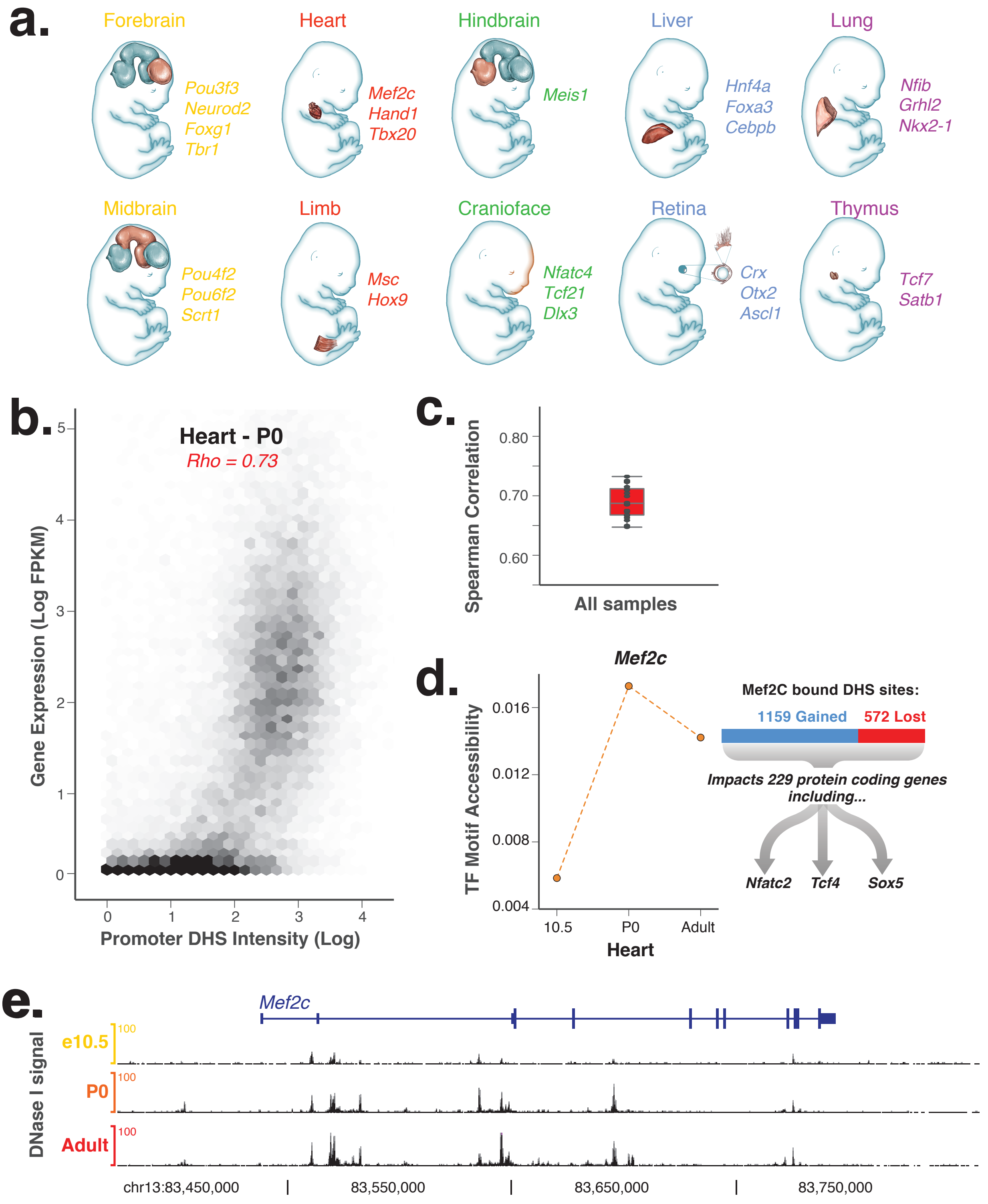


**Figure S2. DHS signal and the regulation of gene expression (A)** Z-score transformed TF motif accessibility robustly recapitulates TF gene expression for a range of tissues, including key lineage regulators such as *Mef2c* in heart, *Nkx2-1* in lung, and *Neurod2* in forebrain. **(B)** Hexbin plot showing the relationship between accessibility near protein coding genes' TSSs (log DHS intensity) and gene expression for heart tissue at P0 (log FPKM). The corresponding correlation value (Spearman's rho) is indicated in red. **(C)** Box-and-whisker plot of the Spearman correlation for all matched RNA and DNaseI-Seq datasets (Box: interquartile range, IQR, Whisker: 1.5*IQR). Superimposed dots indicate the correlation value for a pair of matched RNA and DNaseI-Seq datasets. **(D)** *Mef2c* motif accessibility across heart development (lineplot) and the number of gained/lost DHSs containing *Mef2c* motif between e10.5 and P0 (stacked barplot). By linking DHSs to the nearest gene, 229 protein coding genes are linked to the gained *Mef2c* DHSs. These include other TFs involved in cardiomyocyte biology including *Nfat2c*, and *Tcf4*. **(E)** Genome browser view showing DNaseI cleavage density across the *Mef2c* gene locus for e10.5, P0 and Adult samples.

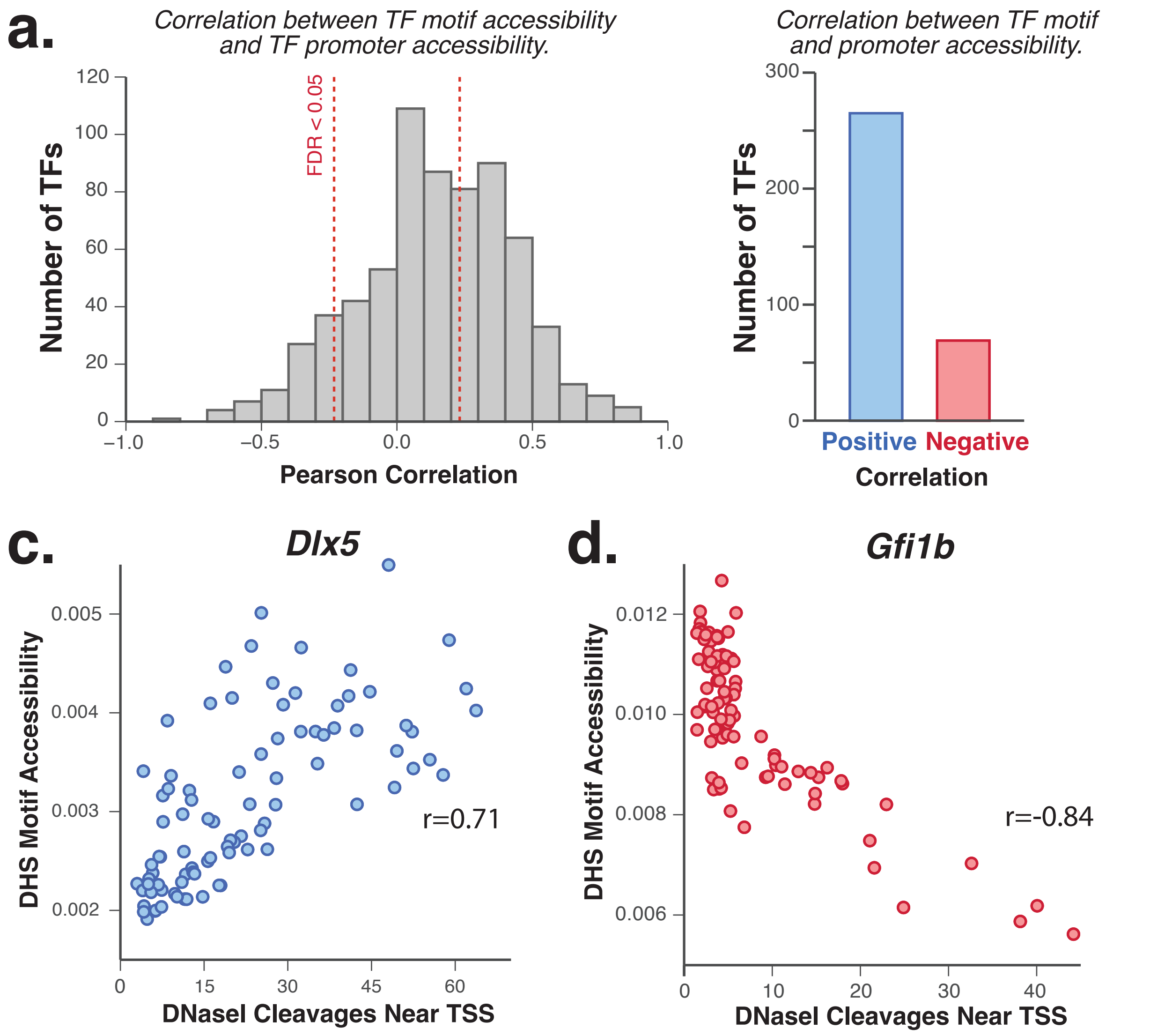


**Figure S3. Correlation of genome-wide motif accessibility with TF promoter accessibility. (A)** Histogram shows Pearson correlation values for the correlation between TF motif accessibility and TF promoter accessibility across samples. **(B)** Number of TFs showing positive or negative correlation between TF motif and promoter accessibility (FDR<0.05). **(C-D)** Scatter-plot indicating the promoter accessibility relative to motif accessibility (y-axis) for 89 samples is plotted for a positively correlated TF (*Dlx5*) and negatively correlated TF (*Gfi1b*).

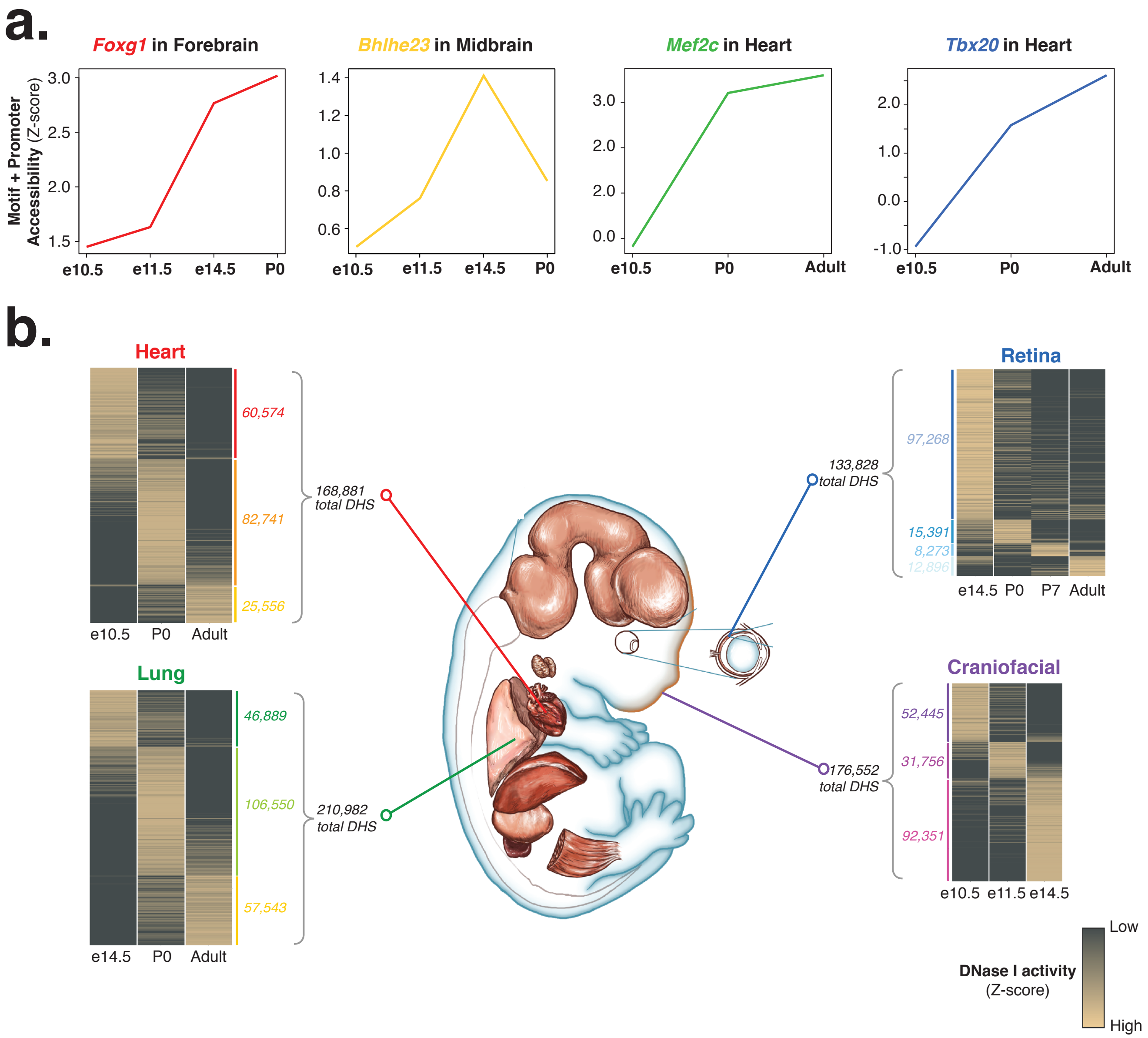


**Figure S4. Dynamic transcription factor motif accessibility across developing mouse tissues. (A)** Lineplots of combined TF motif and promoter accessibility (Z-scores) for selected lineage-specifying TFs across developmental timepoints in relevant tissues. **(B)** Heatmaps indicate variable DNase I accessibility across developmental timepoints in the selected tissues. All variable DHSs are ordered by their time of maximum activity. For each timepoint, the numbers of DHSs exhibiting maximum activity is indicated.

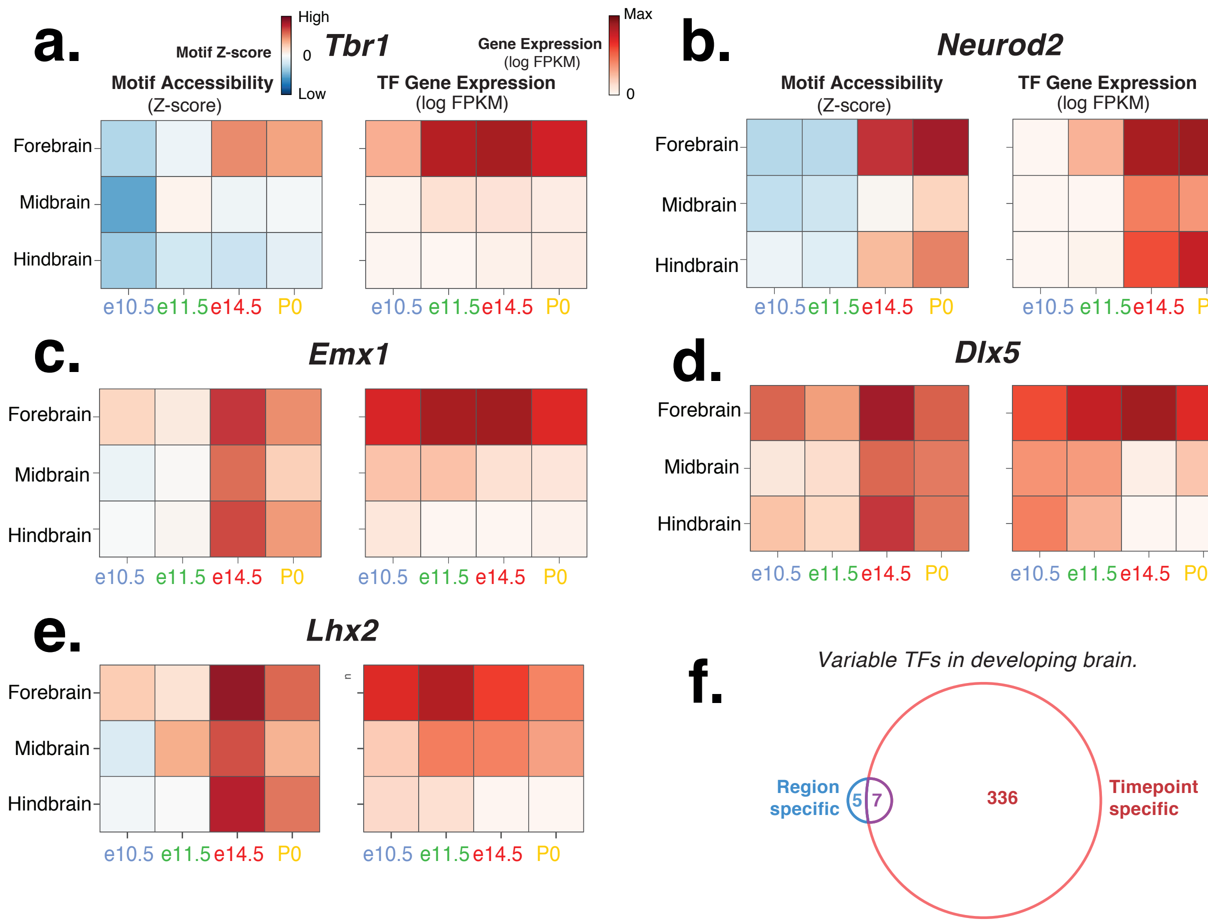


**Figure S5. Differential TF activity during brain development. (A)** RNA gene expression and motif accessibility for *Tbr1*, a TF that was identified as variable across developmental time (F-test BH FDR < 0.1), with activity mostly localised to forebrain. **(B-E)** TF motif accessibility (Z-scores) and RNA-seq expression (FPKM) are plotted as heatmaps for four example TFs, including *Neurod2*, *Emx1*, *Dlx5* and *Lhx2*. Note that some TFs present activity patterns that are more tissue-specific (e.g. *Dlx5*), in contrast with others that are more timepoint-specific TFs (e.g. *Neurod2*). **(F)** Venn diagram showing number of TFs with a combined Z-score that is significantly different across developing brain timepoints or regions (F-Test, FDR<0.1). The majority of TFs in these classes are affected by timepoint rather than tissue.

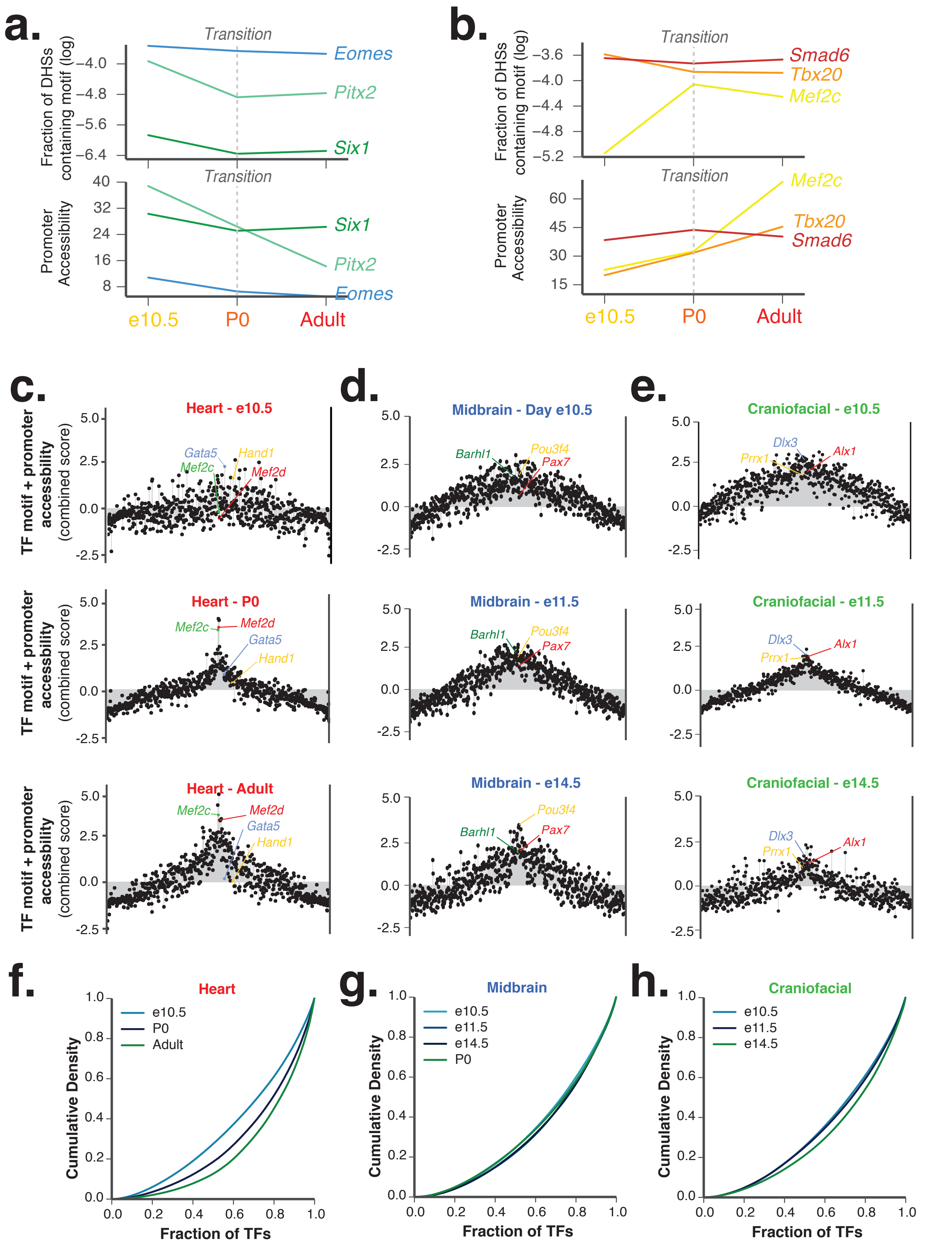


**Figure S6. Dynamic transcription factor motif accessibility across developing mouse tissues. (A)** Lineplots of combined TF motif and promoter accessibility (Z-scores) for selected downregulated TFs across developmental timepoints in heart. **(B)** Same for selected heart lineage-specifying TFs. **(C-E)** TF activity plots indicate variable TF motif accessibility across developmental timepoints in the selected tissues. All variable TFs are ordered randomly from the centre for total motif accesibility change. **(F-H)** Cumulative density plots for heart, midbrain and craniofacial tissue, with each line showing data for a different timepoint.

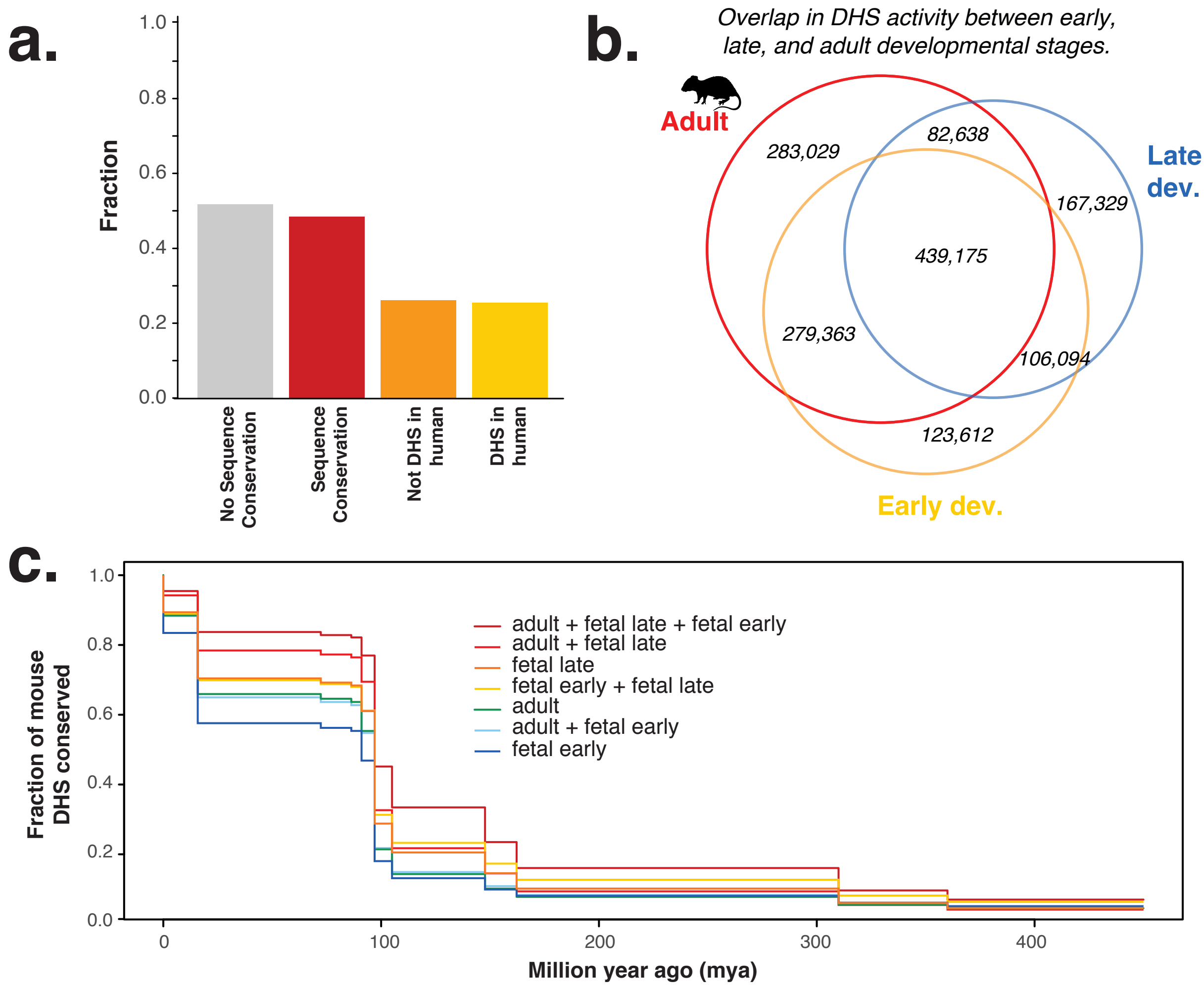


**Figure S7. Conservation of mouse DHS sequence and activity. (A)** Barplot indicates the fraction of mouse DHSs (from all tissues) with no sequence conservation with the human genome, sequence conservation, sequence conservation but not overlapping a human DHS, and sequence conservation and overlapping a human DHS. **(B)** Venn diagram showing the number of ctive DHSs found in different developmental stages (i.e. early fetal, late fetal, adult). **(C)** Lineplot showing the conservation of developmental DHSs present across 500 million years of evolution when compared to all other DHS classes.

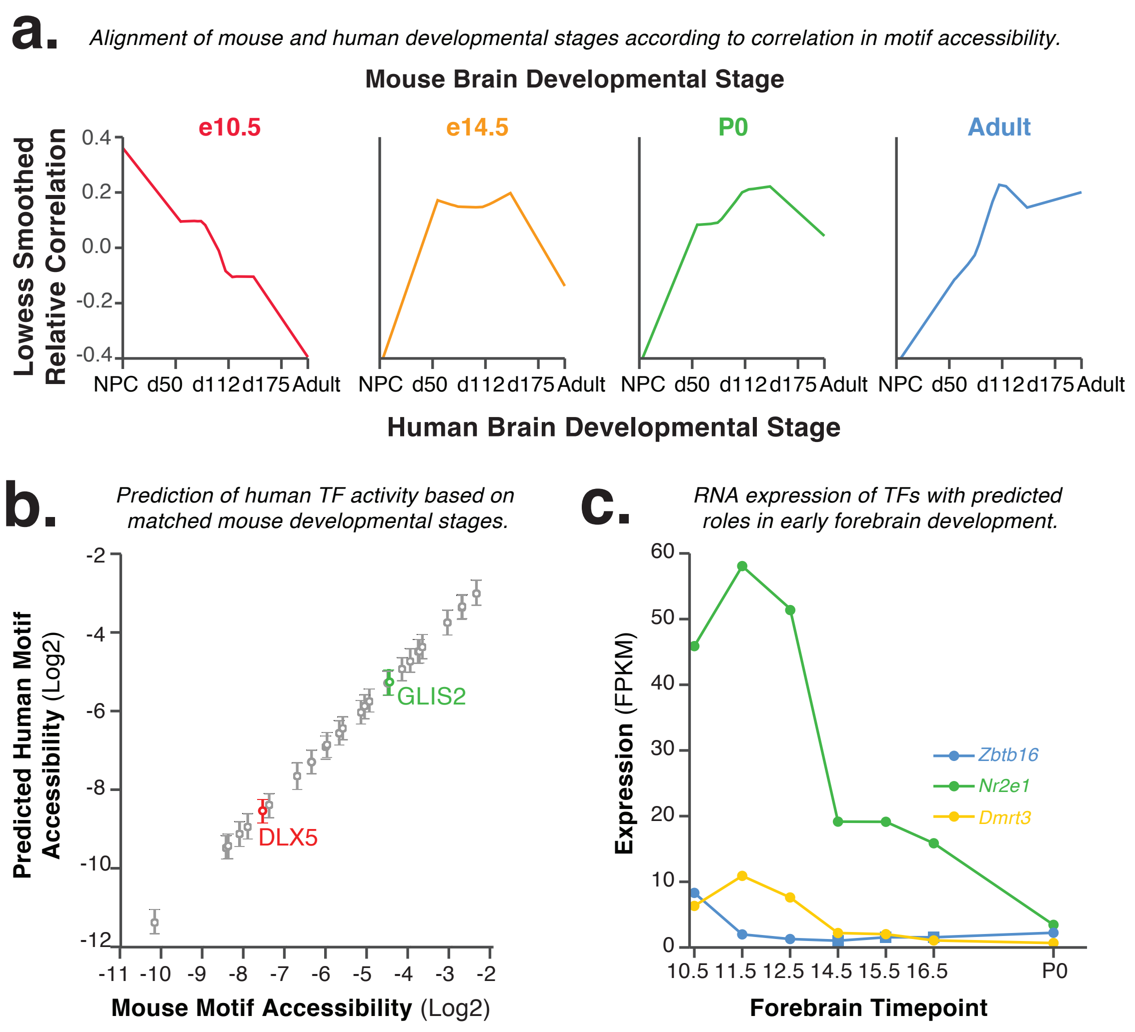


**Figure S8 Alignment between stages of human mouse and human development. (A)** Lowess smoothed relative correlation values between mouse stages in forebrain samples and human brain development. Y-axis is mean-centered to facilitate comparisons, and therefore correlation values are relative to the mean. Notably, the earliest mouse forebrain samples present the highest relative correlation not with human primary samples, but with neural progenitor cells (NPCs). **(B)** Prediction of early human forebrain TF motif accessibility (log of motif accessibility) using mouse TF motif accessibility profiles for early mouse stages. Error bars indicate 50% bootstrapped confidence intervals. For visual clarity, a random subset of all TFs was plotted, in addition to the two examples indicated. **(C)** *Nr2e1*, *Zbtb16*, and *Dmrt3* RNA-seq expression (y-axis) across mouse development (x-axis). The expression of these 3 factors shows a decline during forebrain development, hinting towards a reduction in the activity of these factors in later stages. *Nr2e1*, *Zbtb16*, and *Dmrt3* were predicted to have high activity specific to early (e10.5) forebrain using DNase-seq data.

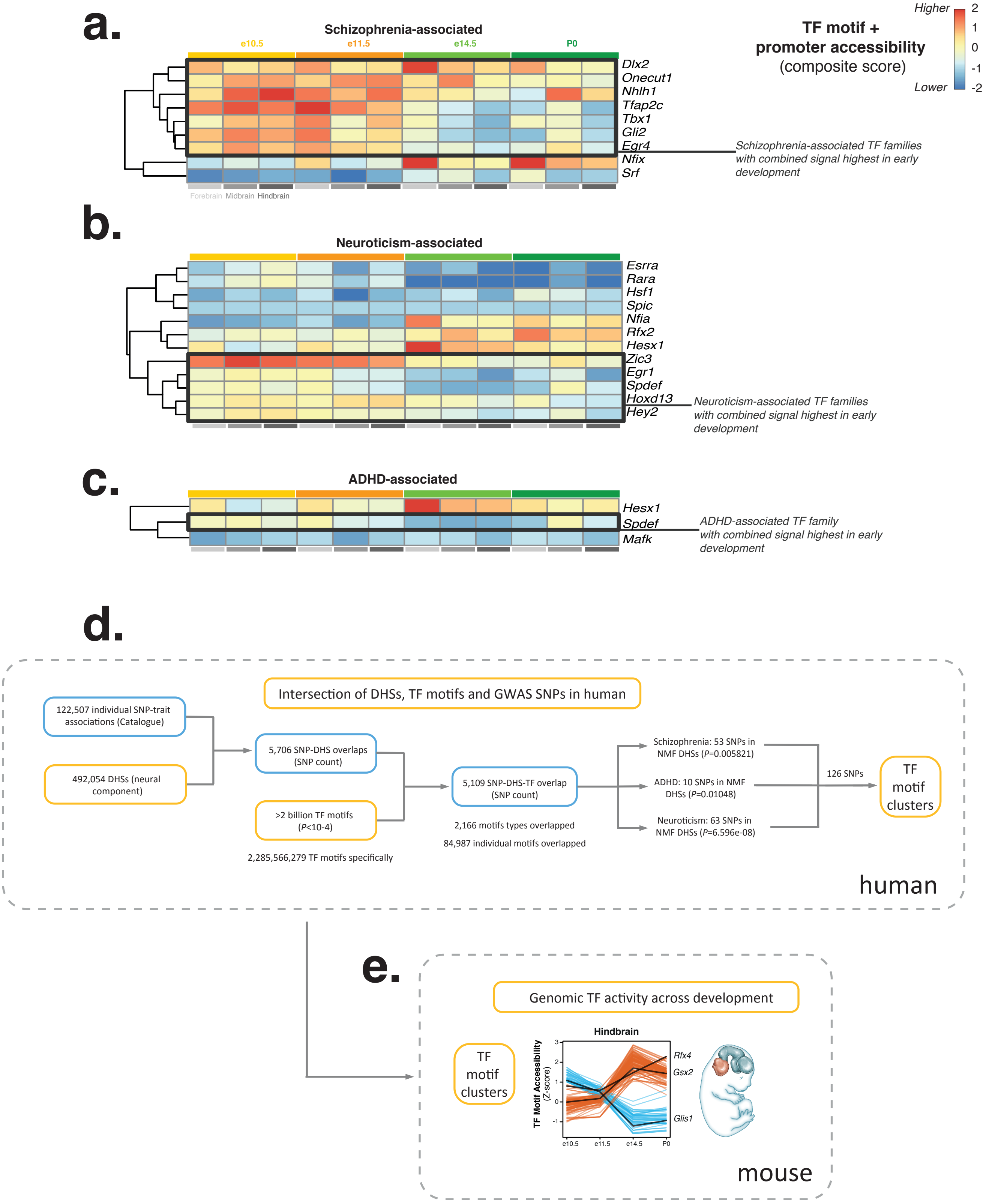


**Figure S9. Early GWAS signal observed for neuronal disease-associated TFs. (A)** Heatmap for the composite score of TF motif and promoter accessibility across brain sections (columns: forebrain, midbrain and hindbrain indicated) and timepoints (3-column sections: e10.5, e11.5, e14.5 and P0). All TFs included contain motifs that overlap schizophrenia-associated GWAS SNPs. Subset of the heatmap comprising early development-associated TF families (grouped through Euclidean clustering), including *Gli2*, *Egr4*, *Tfap2c*, highlighted by weighted black line. **(B)** Same but for neuroticism-associated GWAS SNPs. Early development-associated TF families overlapping neuroticism-associated GWAS SNPs include *Zic3* and *Egr1*. **(C)** Same but for ADHD-associated GWAS SNPs. *Spdef* was identified as an early development-associated TF family overlapping ADHD-associated GWAS SNPs. **(D)** Schematic showing SNP selection and filtering through DHSs and motifs in human, leading to 126 neuronal disease-associated SNPs, which were projected onto TF motifs clusters that are shared between human and mouse. **(E)** Activity for TF motif clusters identified in human was then surveyed across mouse development.
