## Supplemental Methods for "Atlas and developmental dynamics of mouse DNase I hypersensitive sites"

### Materials and Methods

#### Mouse tissue collection

Mouse fetal and newborn tissues were collected at Fred Hutchinson Cancer Research Center in the Groudine laboratory in accordance with the established and reviewed FHCRC IACUC protocol # 1439 to ensure the ethical treatment of the mice. Mouse retinal tissues were collected in the Reh lab at the University of Washington in accordance with established and reviewed protocols (University of Washington IACUC #2448-08) to ensure the ethical treatment of the mice.

#### Fetal and newborn tissue isolation

Timed mating: In the late afternoon a single male was put in a cage with four non-pregnant females and left overnight. The next morning, the male was removed from the cage and the females were checked for plugs, which constitute evidence of mating activity. If a plug was found, that day was considered 0.5 dpc (days post coitum) or e0.5 (post-conception day 0.5, following standard nomenclature, where the e comes from embryonic). Females having a plug were weighed to get a baseline. Plugged females to be used for pre-term embryos were housed together, and plugged females to be used for P0 (postnatal day 0) neonates were housed individually. Mice were periodically weighed to monitor pregnancy. See the table below for approximate expected weight gains.

| **Stage** | **Approximate Weight Gain** |
|---|---|
| E8.5 day | 2-3g |
| E9.5 day | 3-4g |
| E10.5 day | More than 4g |
| E11.5 day | More than 4g |

| | |
|---|---|
| E14.5 day | Pregnancy readily apparent |
| P0 | Pregnancy readily apparent |

#### Staging of embryos

The morning after mating was considered to be day 0.5. Half-day embryos/fetuses (E9.5, E10.5, E11.5 and E14.5) were dissected during the day, preferably in the morning, and full-day embryos/fetuses (E10.0 and E11.0) were dissected in the evening after 7PM. C57BL/6 mice have a gestation of 19.27 days. To be considered P0 (newborn), only pups born in the last 18 hours were included. The chart describes various landmarks of development that were used for staging E9.5 to E14.5 embryos/fetuses. For each sample, embryo tissues from at least 2 pregnant mice were pooled. Any embryo not meeting these staging criteria was excluded from the study.

| **Day:** | **E9.5** | **E10.0** | **E10.5** | **E11.0** | **E11.5** | **E14.5** |
|---|---|---|---|---|---|---|
| **Body rotation** | Complete | Head covers tail tip | Head extends over tail | | Face near HLB | Face more distant from HLB |
| **Face** | | | 2 part first arch visible | | Snout protrudes beyond FB | |
| **Forebrain** | Expands beyond eye | Equal in size to midbrain | | FB largest part of brain | Skin visible over FB | |
| **Eye** | | | Lens of eye becomes visible | | Iris pigmentation visible | Eyelids may be open |
| **Midbrain** | | | Distinct junction with FB and HB | | Skin visible over MB | |
| **Hindbrain** | | | $4^{th}$ ventricle space visible | | Skin visible over ventricles and HB | |
| **Heart** | Ventricle expanded, pale | Atria pink | Atria red, covered by FLB | | Heart less visible (covered by FLB, skin) | Heart even less visible |

| | | | | | | |
|---|---|---|---|---|---|---|
| **Liver** | Small, pale and indistinct | | Pink, small, but more distinct | | Liver prominent under heart | Liver red and easily visible |
| **Intestines** | | Primary loop, internal | | | Starting to protrude into stalk | |
| **Forelimbs and hindlimbs** | | FLB’s close to heart | FLB paddles, smooth, edges visible | FLBs near heart | FLBs at heart, distal indents, HLBs paddles | Fingers separate, long bones of limbs present |
| **Tail** | Next to head | Curls next to head, thinner | Thin, elongated | | | |

*Abbreviations: FB: Forebrain, FLB: forelimb buds, HB: hindbrain, HLB: Hindlimb buds, MB: midbrain*

#### Tissues dissected

Whole embryos: Days E9.5, E10.0, E10.5 and E11.0.

Tissue obtained at day E10.5: Forebrain, midbrain, hindbrain, craniofacial, neural tube, limb, heart, liver and yolk sac.

Tissues obtained at day E11.5: Forebrain, midbrain, hindbrain, craniofacial, neural tube, limb, heart and liver

Tissues obtained at day E14.5: Forebrain, midbrain, hindbrain, craniofacial, limb and lung

Tissues obtained at P0: Forebrain, midbrain, hindbrain, thymus, heart, lung, liver, stomach, kidney

*Mouse retinal dissections.* Whole eyes from E14.5-day embryos were placed in ice-cold Phosphate-buffered saline (PBS). Under a dissecting microscope, sclera and RPE (Retinal Pigment Epithelium) were removed, and forceps were used to separate the lens from the retina. Dissected retinas were pooled for preparation of nuclei.

##### Isolation and cryopreservation of nuclei from tissues

To release the nuclei, tissues were dounced 10 times in 4ml Sucrose buffer (250mM sucrose, 10mM Tris-HC-(pH 7.0, 1mM MgCl2) containing protease inhibitor. Nuclei were filtered through a 40uM mesh filter. DMSO was added to a concentration of 10% and nuclei were frozen in approximately 1 ml aliquots.

#### Cell lines and differentiations/treatments

*Adipocytes***.** 3T3 L1 Pre-adipocytes and day 8 differentiated adipocytes were obtained in collaboration with Sona Kang. Briefly cells 3T3 L1 cell were maintained in Dulbecco's modified Eagle's medium (DMEM) containing 10% calf serum. Post 2-days confluence (adipogenic day 0), cells were induced with DMEM containing, 10% fetal bovine serum (FBS), 5mM methylisobutylxanthine, 1uM dexamethasone and 5ug/ml insulin. On day 2, induction media was removed and replaced with DMEM containing 10% FBS and 5ug/ml insulin. On day 4, insulin containing media was removed and replaced with fresh DMEM containing 10% FBS every other day. Cells differentiated into adipocytes by day 8.

*Motor Neuron-like cells (AR24Q MN1).* These cells were obtained in collaboration with the La Spada lab. This cell line is a fusion of mouse embryonic motor neurons with mouse neuroblastoma cell lines[1]. The cell line was modified to express wild type human androgen receptor, AR24Q[2]. Cells were serum starved for 48 hours, and then treated overnight with testosterone (R1881) at a concentration of 1uM. 48 hours later, cells were harvested and run on a percoll gradient to remove apoptotic cells. Cells were frozen in freezing media: 40% Normosol-R, 40% bovine serum albumin (BSA), 20% dimethyl sulfoxide (DMSO).

*PMTPU cells #10.* These cells were obtained in collaboration with Art Skoultchi. They are a mouse erythroleukemia (MEL) cell line with a Zn+ inducible promoter driving PU-1 expression. Cells were grown in DMEM with 10% FBS and 1 mg/ml G418. They were induced for 24 hour with media containing 125uM ZnCl in order to overexpress PU-1.

*mESC R1 cells.* These cells were obtained in collaboration with the Wysocka lab and were differentiated into epiblast like cells for 12 and 24 hours[3]. Apoptotic cells were removed by percoll gradient before nuclei isolation

#### Isolation of nuclei from primary or cultured cells

Cryopreserved cell were "slow-thawed" by gradual addition of sequentially larger volumes of room temperature PBS supplemented with 1% FBS under constant mixing. This gradual addition prevents damage to cell membranes during thawing by permitting equilibration. Fresh cells were washed with PBS. Whether fresh or frozen, cells were counted using a hemocytometer, centrifuged at 300g and re-suspended in buffer A (15 mM Tris HCl pH 8.0, 15 mM NaCl, 60 mM KCl, 1 mM EDTA pH 8.0, 0.5 mM EGTA pH 8.0) at a concentration of less than $10^7$ cells/ml. In order to isolate nuclei, equal volume of 2x Igepal was added and cells were incubated on ice with occasional mixing. Final Igepal concentration and incubation time are determined for each cell type. For these cell types they are as follows: Preadipocytes: 0.025 % Igepal for 5.5 min; Adipocytes: 0.035% Igepal for 6 min.; AR24Q MN1: 0.7 % Igepal for 3 min; PMTPU cells #10: 0.015% Igepal for 6 min and mESC R1, 0.025% Igepal for 5 min.

#### DNase I treatment and sequencing

Nuclei are counted, pelleted at 500g and resuspended in 37°C DNase I digestion buffer (13.5 mM Tris-HCL pH 8.0, 87 mM NaCl, 54 mM KCl, 6 mM CaCl2, 0.9 mM EDTA, 0.45 mM EGTA, 0.45 mM spermidine), with concentrations of DNase I (Sigma-Aldrich) ranging from 0 to 100U/ml. Nuclei were digested at 37°C for 3 minutes and the reaction was stopped with 1 volume of 2X stop buffer (50 mM Tris-HCl pH 8.0, 100 mM NaCl, 0.1% SDS, 100 mM EDTA pH 8.0, 1 mM spermidine, 0.3 mM spermine). RNAse was added and the sample was incubated at 37°C for 1 hour. Proteinase K was then added and the sample was incubated at 55°C for an additional hour. Small DNA fragments were isolated either on a sucrose gradient or by PEG fractionation. Libraries were constructed by ligating adapters ends of purified fragments and the libraries were sequenced on an Illumina Hi-Seq 2000.

#### Sequence alignment of DNase-seq experiments

To generate the mouse DNase-seq reads to the genome we applied the same mapping procedure as described in the recent ENCODE manuscript on indexing accessible DNA elements in the human genome[4]. Data processing of DNase sequencing FASTQ uses the ENCODE DCC DNase-DHS pipeline, version 2, paired-end (ENCPL202DNS). Briefly, reads were trimmed of adapter matched sequence and subsequently aligned using BWA, version 0.7.12. to the mouse genome reference mm10-minimal (ENCSR425FOI). Filtered, quality mapped, reads from all sequencing runs of a given library were merged and subsequently processed for enrichment calls using hotspot2. Further processing and QC evaluation was performed in accordance with the pipeline specification: https://www.encodeproject.org/data-standards/dnase-seq/

#### Annotation of DHSs

The Hotspot2 pipeline for processing DNase I sequencing data takes in Illumina sequencing reads, analyzes the data, and returns genomic regions with statistically significant enrichments, or "hotspots" of cleavage activity via the DNase I experiment. The enrichment caller is available, stand alone, on Github at https://github.com/Altius/hotspot2. This tool leverages an enzymatic DNA cleavage model to measure enrichment over background and identify any resultant DNase I Hypersensitive Sites (DHSs) with resultant confidence values. Alignment files are input into the pipeline and processed over uniquely mappable regions of the genome, the resulting output is a resource used to derive confidence thresholded peak regions (DHSs) in bed format. For one midbrain sample, we used replicated hotspot peaks from two technical replicates, DS33935 and DS30060, in place of the peaks called from a single sample.

#### Atlas generation and annotation

Briefly, we used DHS position, height, width and local distribution across 197 samples to create an atlas or index of mouse regulatory DNA (available in BED format as **Supplementary File 1**). Strong (FDR < 0.1%) variable-width peaks were called using hotspot2 (Rynes et al., in preparation). An atlas is created from these per-sample peaks by calling consensus DHS positions across 197 samples, incorporating both element position and dispersion across tissues and time points (https://github.com/Altius/Index)[4]. ENCODE DCC library IDs for these samples are listed in **Supplementary File 2**.

#### Annotation of DHS tissue/cell selectivity

To annotate the cell type-specificity of DHSs, atlas elements were defined as active in a sample if the overlap between atlas element and sample DHS was >= 25% of the element width. In all cases DHSs were called using both sample replicates except for a reduced set of samples where only one replicate was available. Constitutive DHSs were defined as DHSs detected in greater than 75% of samples. Timepoint-specific DHSs were defined as the subset of tissue-specific DHSs active at only one developmental time point.

#### Reproducibility between biological replicates

DNase-seq reads from matched tissue samples from different animals were highly correlated (mean log [read count] pairwise correlation of r=0.80 vs. r=0.55 between un-matched tissues; **Figure S1A**).

#### K-means clustering

The number of DNase I cleavages falling within each atlas element was counted, and data were quantile normalized as a standard procedure to correct for library quality differences. Replicates for a given sample, where available, were averaged. K-means clustering was performed on the averaged data using sklearn[5]. The number of clusters (35) was chosen based on the rate of change of the total squared distance between data points and their assigned cluster center using the elbow method[6,7].

#### TF motif analysis

TF motifs were identified by scanning mm10 using the FIMO tool[8] from the MEME suite[9] (motif match $P$<$10^{-5}$) with position weight matrices from the JASPAR[10], TRANSFAC[11], and UNIPROBE[12] databases and SELEX data[13].

#### TF activity measures

We used two measures of TF activity, one focused on the TF gene itself and one focused on the genome-wide landscape of its cognate recognition sequence.

*TF promoter activity.* For TF gene activity, we exploited the fact that quantitative DNA accessibility of the promoter region of a given gene is very tightly correlated with transcriptional output (average r > 0.93)[14,15,16]. We therefore computed this parameter for the interval spanning +/-5kb of the promoter for all annotated transcription factor genes in the mouse genome (n=638). For the subset of TFs that share the same or highly similar cognate motifs, the specific family members with activated promoters were annotated.

*Cognate TF landscape / genome-wide TF motif accessibility*. We used the measure defined in Vierstra et al.[17], the proportion of DHSs within a given cell/tissue type that contain at least one cognate motif for a given TF. We quantified this fraction for each TF and its cognate motif for each tissue/cell sample. This parameter was found to be the most conserved regulatory feature between human and mouse[17]. TFs play a direct role in mediating DNA accessibility at regulatory DNA. When the cognate of a given TF is present within accessible regulatory DNA (i.e. a DHS), the probability of occupancy by that TF is very high and directly parallels orthogonal measures of occupancy such as a ChIP-seq[18,17,19].

*Composite measure of TF activity.* We Z-score transformed both the TF promoter accessibility and TF landscape scores, and added the Z-scores to obtain a composite measure of TF activity. While chromatin accessibility around most TFs' TSSs was positively correlated with motif accessibility, a subset of TFs showed a strong negative correlation between promoter accessibility and motif activity. These TFs included known repressors including Gfi1 and Gfi1b as well as Zbtb16. For these TFs we reversed the sign of the motif Z-score when calculating the combined TF activity since lower motif accessibility in DHSs likely indicates higher activity of the TF.

#### Mouse vs. Human TF comparisons

We exploited the high correlation between *trans* regulatory activity in equivalent human and mouse tissues to predict tissue-specific human TF activity based on mouse DHS data in tissues and developmental timepoints that are difficult to obtain. First, to quantify TF activity, we measured the fraction of DHSs containing the TFs' motifs in each cell type. We then Z-score transformed the motif fractions (separately in human and mouse datasets) to score tissue-specific motif activity. We identified 22 closely matched samples between the human and mouse datasets, and using the matched samples, we fit a linear mixed model to predict human motif activity from the mouse data (model: Log(Human.Motif_Density) ~ Log(Mouse.Motif_Density) + ( 1 | TF ) + ( Log(Mouse.Motif_Density) | Cell_Type ) fit using the R package lme4). From the model that was fit using the 22 closely matched samples, we then predicted tissue human motif activities (Z-scores) for samples where the human data is not available from the mouse data and bootstrapped 50% prediction intervals (holding the random effects constant) to estimate the uncertainty in the prediction.

#### Time point-specific DHSs and TFs

To identify developmental time-point specific DHSs we quantified the activity of the DHS by the number of cleavages in each element, normalized the data across samples using DESeq's normalization strategy restricted to constitutive peaks, and log-transformed the normalized counts (log[cleavages + 1]). We identified time-point variable DHSs as replicated DHSs that were not called at all timepoints across development. We then ordered the time-point variable DHSs first on the time of their maximum signal and secondarily by a time average of their signal.

To identify the TFs underlying the time point-specific regulatory activity, we identified TFs whose genome-wide motif accessibility varied the most across the timepoints (heatmaps include all TFs with motif activity >1 Z-score and a difference of >1.5 between the minimum and maximum activity).

#### Brain ANOVA

In brain, where we had the most comprehensive mapping of chromatin accessibility, we performed a regression on each TF using developmental time as a categorical variable (Combined-Zscore ~ Time + Region + Region:Time). We compared the full model to a reduced model (F-test, Combined-Zscore ~ 1) to identify TFs that varied over the time course or brain regions. We identified TFs that differed across regions by comparing to the reduced model Combined-Zscore ~ Time and the reduced model Combined-Zscore ~ Region for TFs that vary across development stages. We performed a similar procedure for each DHS (full model: Log(DNaseI Cleavage + 1) ~ Time + Region + Region:Time) to identify DHSs with significantly varying accessibility.

#### Gini analysis

Briefly, an F-test comparing the full model that includes tissue, time, and the interaction (gini~tissue + time + tissue:time) to the reduced model in which just tissue was included as an explanatory variable (between gini~tissue) showed a highly significant p-value of 5.11e-08.

#### GWAS analysis

GWAS analysis was performed as a three-step process. First, we identified the human TF motifs that co-located with neural component DHS-overlapping GWAS SNPs[4]. Second, we analysed the activity of the corresponding mouse TFs across all mouse stages. Third, we identified a subset of TFs that present a high signal at the earliest mouse developmental samples (e10.5-e11.5, corresponding to stages for which equivalent human samples are impossible to obtain).

Focusing on brain, we took GWAS SNPs overlapping human neural component DHSs, and overlapped them with human TF motifs. We started with 122,507 individual SNP-trait associations from the GWAS catalogue (https://www.ebi.ac.uk/gwas/, downloaded 2-September-2019). We used SNP-trait associations because one SNP can be associated with several traits. This number was then reduced to 5,706 SNP-trait associations that overlapped at least one of the 492,054 human neural component DHSs (generated as described in the ENCODE manuscript on indexing accessible DNA elements in the human genome)[4]. Of these SNPs, 5,109 overlapped TF motif matches (motif matches called at p-value < 1e-04, using FIMO[8]). We then focused on the activity of the TFs for 4 neuronal diseases that presented enrichment in the neural component DHSs: schizophrenia, depressive symptoms, attention deficit hyperactivity disorder (ADHD) and neuroticism. For these diseases, we focused on the TFs that presented the highest overlap from the final list, aggregating different TF motif matches by TF gene names. We then plotted and clustered the genome-wide motif accessibility for these TFs in mouse brain samples across e10.5-P0 to identify TFs with a highly accessible DNA landscape at e10.5-e11.5.

#### RNA-seq analysis

We downloaded processed data (FPKMs) for 166 RNA-seq samples across 17 developing mouse tissues from the ENCODE consortium webpage (https://www.encodeproject.org/, accessed 02/13/2017). Samples included were mapped to mm10. Only primary tissue samples were selected. All tissues included at least one sample taken during mouse fetal development (up to P0). Timepoints comprised e10.5-postnatal week 8. A supplementary file including the FPKMs used is available (**Supplementary Table S4**, tab-separated values, matrix columns: ENSEMBL gene ID, ENSEMBL transcript ID, and sample-time point name).
